# Genetic manipulation of a giant virus-associated virophage

**DOI:** 10.64898/2026.06.16.732491

**Authors:** Jingjie Chen, Hiroyuki Ogata, Hiroyuki Hikida

**Affiliations:** Chemical Life Science, Institute for Chemical Research, Kyoto University, Uji, Kyoto 611-0011, Japan; Department of Virology I, National Institute of Infectious Diseases, Japan Institute for Health Security, 1-23-1 Toyama, Shinjuku, Tokyo 162-8640, Japan; Research Center for Biosafety, Laboratory Animal and Pathogen Bank, National Institute of Infectious Diseases, Japan Institute for Health Security, 1-23-1 Toyama, Shinjuku, Tokyo 162-8640, Japan

**Keywords:** Virophage, Giant virus, Reverse genetics, Circular polymerase extension reaction

## Abstract

Virophages are double-stranded DNA viruses that hyperparasitize giant viruses infecting unicellular eukaryotes. Parasitization by virophages often reduces the replication of giant viruses, thereby modulating microbial communities in the environment. However, the molecular mechanisms underlying the tripartite relationship are largely unknown due to methodological limitations. In the present study, we developed a reverse-genetics system for a Sputnik virophage that parasitizes the amoeba-infecting giant virus, mimivirus. We demonstrated that transfection of genomic DNA could recover infectious virophage particles. Transfection of genomic DNA synthesized by circular polymerase extension reaction (CPER) also resulted in the recovery of infectious viruses. As a proof of concept, we successfully modified two Sputnik genes by transfecting CPER-assembled mutant genomic DNA. Collectively, our reverse-genetics system provides a framework for assessing the functional importance of Sputnik genes and should facilitate future genetic studies of virophages.

**Significance statement:** Virophages are viruses that hyperparasitize giant viruses, which infect unicellular eukaryotes and have extremely large particles and genomes. Giant viruses modulate microbial communities not only by killing their hosts but also by altering host cellular functions. Virophages modulate the replication of giant viruses, thereby driving ecosystem dynamics. Previous studies have demonstrated their widespread distribution through isolation and metagenomic analyses. However, the functions of most virophage genes remain unknown. Due to the lack of genetic tools, the molecular mechanisms underlying the interactions between virophages and giant viruses remain largely elusive. Here, we established a virophage reverse-genetics system based on circular polymerase extension reaction. Our results demonstrate that the system can dissect virophage gene functions and will accelerate virophage genetics.

## Introduction

Virophages are double-stranded DNA viruses that parasitize giant viruses, which infect unicellular eukaryotes. Previous studies have identified diverse combinations of virophages, giant viruses, and eukaryotic organisms in various environments (1–3). The presence of virophages often reduces the replication of host giant viruses, thereby modulating microbial communities. Recent studies have characterized the transcriptional landscape of model virophage culture systems (4, 5). However, molecular interactions in these tripartite systems remain largely elusive due to the lack of reverse-genetics systems applicable to virophages.

Circular polymerase extension reaction (CPER) is a modified method of polymerase chain reaction (PCR) for generating circular DNA from overlapping PCR fragments. This method skips bacterial cloning, enabling the rapid construction of recombinant genomes free from limitations imposed by bacterial culture (6). These advantages are particularly valuable for viral reverse-genetics systems. Accordingly, CPER has been widely used in reverse-genetics systems for diverse viruses (7–10).

In the present study, we confirmed that genomic DNA is sufficient to recover infectious particles of a Sputnik virophage, which parasitizes the amoeba-infecting giant virus mimivirus. We then developed a CPER-based reverse genetics system for the Sputnik virophage and demonstrated its utility for virophage genetics.

## Results and Discussion

### Recovery of a replication-competent Sputnik virophage from its genomic DNA

Viral reverse genetics systems often recover infectious particles from their genomic DNA or RNA, allowing viral sequences to be readily modified in bacteria or by PCR. Because some virophages are integrated into the genomes of eukaryotes or their host giant viruses (11, 12), we reasoned that genomic DNA might be sufficient to recover infectious virophages. We extracted intact virophage DNA from Sputnik 3 particles, then transfected the DNA into amoeba cells, and inoculated the cells with Acanthamoeba polyphaga mimivirus (APMV) (Fig. 1A). We successfully detected Sputnik genomic DNA from the supernatant, the amount of which increased after passaging the supernatant (Fig. 1B). We also found that the supernatant of passage one (P1) contained Sputnik particles (Fig. 1C). Collectively, these results indicate that the transfection of Sputnik genomic DNA can recover replication-competent Sputnik.

**Fig 1.**
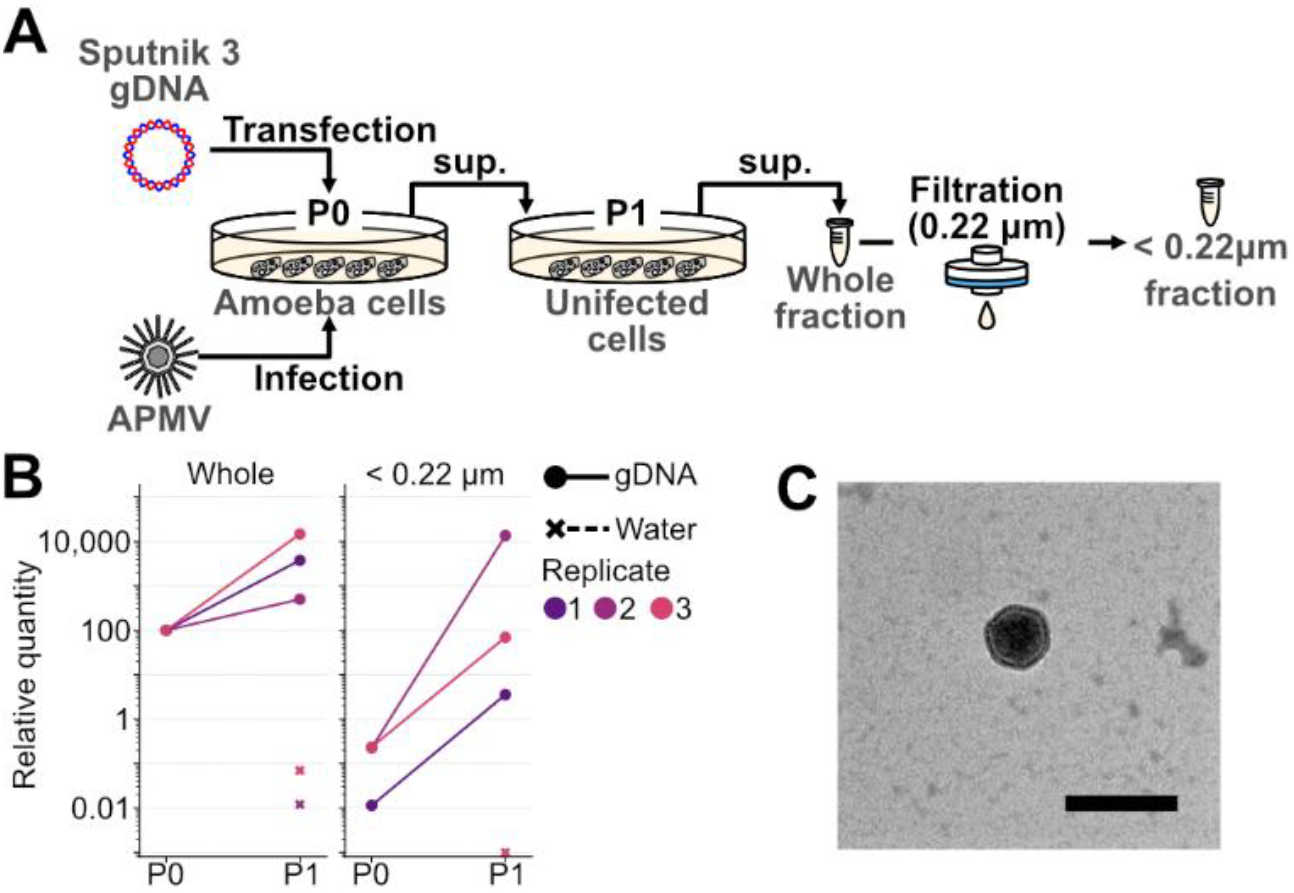
(A) Schematic illustration of Sputnik 3 recovery following transfection of its genomic DNA. (B) Relative quantity of the genomic DNA of Sputnik. Marker and line styles indicate genomic DNA transfection and water treatment (negative control). Each color represents a replicate. Sputnik genomic DNA levels are expressed relative to the total amount in the genomic DNA-transfected well, which was set to 100. Dots were connected by lines when genomic DNA was detected in both P0 and P1; otherwise, only individual dots are shown. (C) Negative staining image of supernatant from the genomic DNA-transfected well at P1. Bar = 100 nm.

### CPER successfully modified the Sputnik genome

Because CPER facilitates viral reverse genetics, we used it to assemble Sputnik genomic DNA and establish a reverse-genetics system for this virophage. We divided the Sputnik genome into four fragments with overlapping sequences (∼30–35 bp) (Fig. 2A). One overlap lies within the coding sequence of the major capsid protein (*mcp*), which is presumably essential. Thus, we designed four types of constructs for *mcp*: wild type (WT), synonymous mutation (SYN), insertion of a premature stop codon (STOP), and a frameshift mutation (FS) (Fig. 2B). Each construct was transfected, and APMV was inoculated as described above. We detected Sputnik virophage in the resultant P1 supernatant from the WT, SYN, and FS constructs (Fig. 2C). In WT and SYN, Sanger sequencing confirmed the designed sequences, demonstrating that genomic DNA constructed by CPER yields viable Sputnik particles and that its sequence can be edited (Fig. 2C). The STOP construct did not yield viable Sputnik particles. The FS sequence in the P1 supernatant was wild type rather than the intended mutant sequence (Fig. 2C). These results indicate that the loss-of-function mutations in *mcp* were lethal, as expected, and that CPER-based genomic DNA construction can assess the functional importance of Sputnik genes.

**Fig 2.**
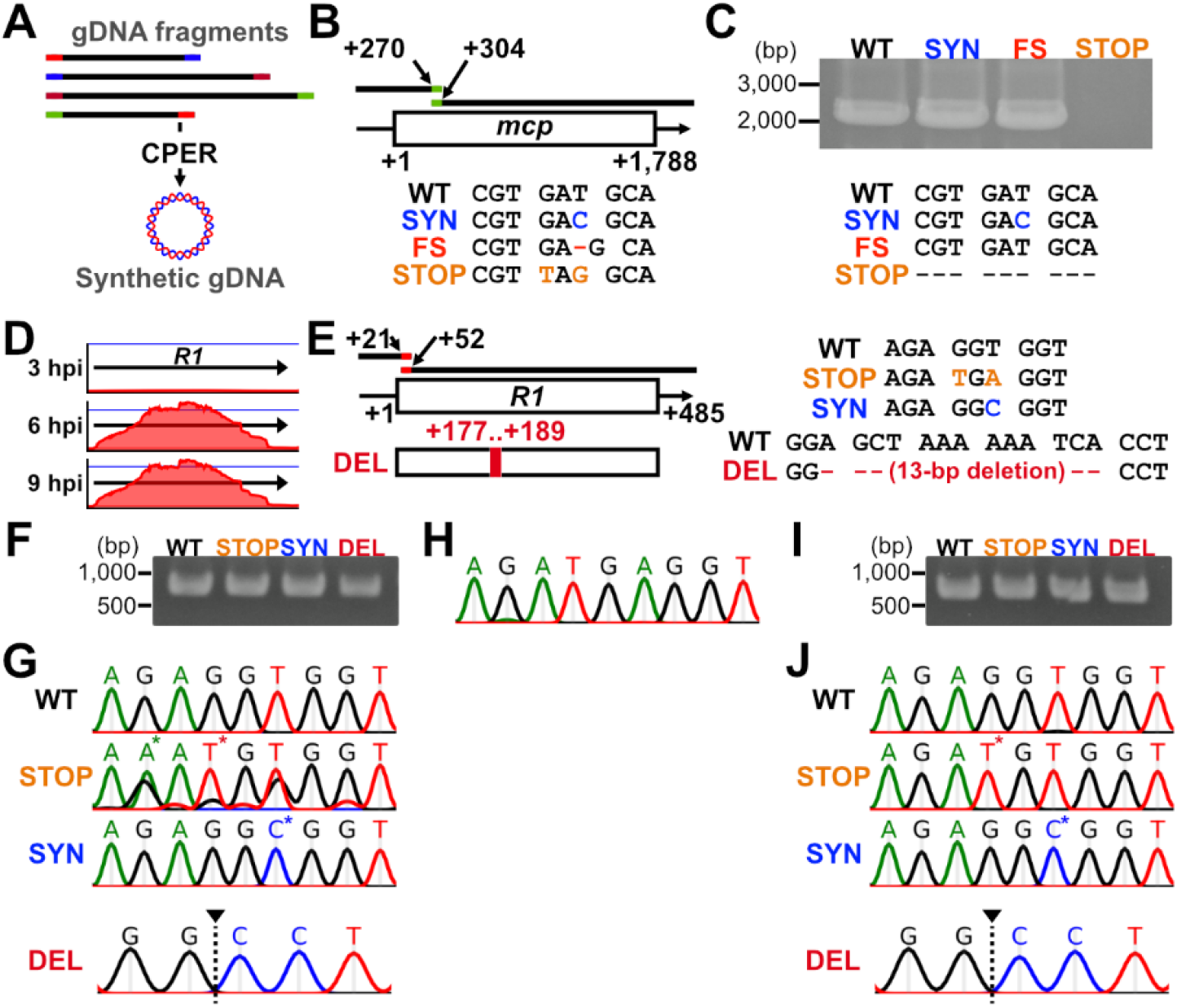
(A) Schematic illustration of CPER. Each bar represents a fragment of Sputnik genomic DNA, and colored regions at the ends indicate overlapping sequences. (B) Schematic illustration of mutations introduced in the *mcp* gene. (C) PCR amplification of part of the *mcp* gene and corresponding partial Sanger sequencing results from the amplicons. DNA was extracted from the P1 supernatant. (D) Transcription of the *R1* gene. RNA-sequencing data from a previous transcriptome analysis (4) are shown at single-nucleotide resolution. Arrows indicate the coding sequence of *R1*. Blue lines indicate a read depth of ×300,000. (E) Schematic illustration of mutations introduced into the *R1* gene. (F) PCR amplification of the partial *R1* gene in replicate 1. (G) Partial Sanger sequencing results from the amplicons in replicate 1. Mutations and deletions are indicated by asterisks and black arrowheads, respectively. (H) Partial Sanger sequencing of the CPER product for the STOP construct in replicate 1. (I) PCR amplification and (J) Sanger sequencing results in replicate 2. (F-J) DNA was extracted from the P1 supernatant.

Another overlap lies in the *R1* coding sequence, a late gene encoding a hypothetical protein (Fig. 2D). As we failed to introduce a frameshift mutation in *mcp*, we designed an alternative loss-of-function construct for *R1* containing a larger deletion expected to disrupt gene function (DEL), together with WT, SYN, and STOP constructs (Fig. 2E). We successfully recovered the Sputnik virophage from the WT construct as well as from the mutant constructs. However, Sanger sequencing indicated that the STOP population was not clonal (Fig. 2F, G). We confirmed that the CPER product carried the intended mutation, suggesting that this heterogeneity arose or was amplified during passaging (Fig. 2H). Although we attempted to make another replicate, one of the two intended substitutions in STOP was not introduced, resulting in a cysteine codon (TGT) instead of a stop codon (Fig. 2I, J). These findings indicate that CPER-based genome construction can be unstable depending on the site and mutation. Nevertheless, we successfully recovered loss-of-function mutants (i.e., a frame-shift deletion in replicates 2 and 3), which suggests that the *R1* gene is dispensable for viral replication.

In the present study, we established a CPER-based reverse-genetics system for the Sputnik virophage, providing a platform for future genetic studies in virophages. At present, the success rate of virophage genome editing varies by target site and mutation. Although previous studies have demonstrated the accuracy of CPER (7, 9), its fidelity is lower than that of bacterium-based methods, such as bacterial artificial chromosome-based systems (13). This variability may reflect PCR-derived errors and could potentially be mitigated by using higher-fidelity polymerases or incorporating bacterium-based cloning steps.

## Materials and Methods

Virophage genomic DNA was prepared from Sputnik 3 particles by phenol-chloroform extraction. Genomic fragments were prepared using KOD One PCR Master Mix (TOYOBO). CPER was performed using PrimeSTAR GXL DNA Polymerase (TAKARA). Quantification of Sputnik genomic DNA and transfection were performed as described in previous studies (4, 14). Detailed descriptions are provided in the SI Appendix.

## Acknowledgements

We thank Prof. Bernard La Scola and Ms. Lina Barrassi for kindly providing APMV and Sputnik 3. The electron microscopy study was supported by the Division of Electron Microscopic Study at the Center for Anatomical Studies, Graduate School of Medicine, Kyoto University.

This study was supported by Japan Society for the Promotion of Science KAKENHI grants 25K18450, 22K15175 to H.H. and 22H00384 to H.O., and by Japan Science and Technology Agency ACT-X Grant JPMJAX22BI to H.H.

