## Supplementary material for "Genetic manipulation of a giant virus-associated virophage": SI Appendix

This file includes:

- Extended Methods
- SI References

### Extended Methods

#### Cells and viruses

*Acanthamoeba castellanii* (Douglas) Page, strain Neff (ATCC 30010), was maintained in peptone-yeast extract-glucose (PYG) medium at 28°C. Acanthamoeba polyphaga mimivirus (APMV) and Sputnik virophage 3 (Sputnik) were used as prototypes of mimiviruses and virophages, respectively.

#### Genomic DNA extraction

Culture supernatant of *A. castellanii* inoculated with Sputnik and APMV was collected into a 50 mL conical tube and centrifuged at 2,000 rpm (Thermo Scientific, Sorvall ST 8FR) for 5 min at 4°C. The supernatant was filtered through a 0.22 µm filter, then centrifuged at 9,000 rpm for 1 h at 4°C. The pellet was resuspended in 1 mL of phosphate-buffered saline (PBS) and centrifuged at 9,000 rpm for 1 h at 4°C. The pellet was resuspended with Tris-EDTA buffer at pH 8.0. The resuspension was mixed with 5% sodium dodecyl sulfate, 2 mg/mL Proteinase K, and 200 µg/mL RNase A, then incubated overnight at 56°C. Genomic DNA was extracted using equal volumes of phenol and chloroform, each extracted twice, followed by ethanol precipitation.

#### Transfection of genomic DNA

Transfection was performed as described in a previous study (1). Briefly, $5\times10^5$ amoeba cells were seeded into 35-mm Nunc™ EasYDishes or Nunc™ Cell-Culture Treated 6-well dishes (Thermo Fisher Scientific), incubated for 15 min at room temperature, and the medium was replaced with 2 mL of fresh PYG medium. One µg of Sputnik 3 genomic DNA or 20 µL of circular polymerase chain reaction (CPER) product was mixed with 10 µL of Polyfect (Qiagen) reagent in 100 µL of PBS and incubated for 10 min at room temperature. After incubation, 600 µL of PYG medium was added, and the mixture was gently pipetted two to three times before being added to the amoeba cultures. Immediately after the transfection, APMV was inoculated at a multiplicity of infection (MOI) of 10. The dishes were incubated at 30°C.

The culture supernatant was collected at 3 days post-transfection and defined as the Passage 0 (P0) supernatant. The supernatant was centrifuged at 2,000 rpm for 5 min at 20°C and filtered through a 1.2 µm filter. Ten µL of the P0 supernatant was added to amoeba cultures prepared by seeding $5\times10^5$ cells into 35-mm Nunc™ EasYDishes or Nunc™ Cell-Culture Treated 6-well dishes. The dishes were incubated at 30°C for 3 days, and the supernatant was collected as described above and defined as P1 supernatant.

#### Quantification of Sputnik genomic DNA

Viral particles in the culture supernatant were collected into microcentrifuge tubes and centrifuged at 2,000 rpm for 5 min at 4 °C to remove cellular debris. The resulting supernatant was defined as the whole fraction. An aliquot of the supernatant was filtered through a 0.22 μm filter, and the resulting fraction was defined as the Sputnik fraction. The supernatant was further centrifuged at 15,000 rpm for 1 h at 4°C. The pellets were resuspended in 45 μL of 50 mM NaOH and incubated at 95 °C for 10 min. After incubation, 5 μL of 1 M Tris-HCl buffer (pH 8.0) was added, and the mixture was diluted with 450 μL of Tris-EDTA buffer (pH 8.0). The resulting viral DNA solution was diluted with ultrapure water and subjected to quantitative real-time polymerase chain reaction (qPCR) as previously described (2). Briefly, the PCR solution was prepared using KAPA SYBR qPCR mix (KAPA BIOSYSTEMS) with 0.2 μM of each primer. The PCR was performed as follows: initial denaturation at 95°C for 3 min, followed by 40 cycles of 95°C for 15 s, 60°C for 20 s, and 72°C for 10 s. After amplification, a melt-curve analysis was performed by increasing the temperature from 60 °C to 98 °C in 0.3 °C increments. The primers used for qPCR are listed below.

Sptk3_mcp_F 5′-GAGATGCTGATGGAGCCAAT-3′

Sptk3_mcp_R 5′-CATCCCACAAGAAAGGAGGA-3′

#### Negative staining

Negative staining of viral particles was performed as described previously (3). Briefly, cultured viruses were collected from P1 supernatant by centrifugation at 15,000 rpm for 1 h at 4°C, and the pellet was fixed with 1% glutaraldehyde. The fixed samples were stained with uranyl acetate and observed by an H-7650 transmission electron microscope (Hitachi).

#### Primer design for CPER

The Sputnik 3 genome was divided into four PCR fragments of approximately 3–5 kbp each, with 30–35 bp of overlap at both ends. Each overlap sequence was designed to have an approximate melting temperature of 70°C. The primers used for amplifying the fragments are listed below.

Sptk3_CPER_fragment_1_F 5′-TGGAACGCCAATGAGAGGTGG-3′

Sptk3_CPER_fragment_1_R 5′-CCAGCTTGACCCTTGAGGCC-3′

Sptk3_CPER_fragment_2_F 5′-AAACTGGAAATGATGGCCTCAAGG-3′

Sptk3_CPER_fragment_2_R 5′-GACGTTTGATTTCGGCATACTGAAGAG-3′

Sptk3_CPER_fragment_3_F 5′-CTTCGGCACTCTCTTCAGTATGC-3′

Sptk3_CPER_fragment_3_R 5′-GGAATGCACGGAATGCATCACG-3′

| | |
|---|---|
| Sptk3_CPER_fragment_4_F | 5′-ACTTCAACCTGGTCGTGATGC-3′ |
| Sptk3_CPER_fragment_4_R | 5′-CACCACAACAACCACCTCTCATTG-3′ |

**Primer design for mutagenesis**

For targeted mutagenesis, mutations were introduced into overlapping sequences of two primers. Mutations in *mcp* and *R1* were introduced using the overlap between fragments 3 and 4 and between fragments 4 and 1, respectively. For deletion mutants, primers were designed to amplify the upstream and downstream regions flanking the target deletion site, with complementary overlap sequences added for overlap extension. The primers used to amplify the fragments by PCR are listed below.

| | |
|---|---|
| Sptk3_CPER_fragment_4_stop_F (*mcp*) | 5′-ACTTCAACCTGGTCGTtagGC-3′ |
| Sptk3_CPER_fragment_3_stop_R (*mcp*) | 5′-GGAATGCACGGAATGCctaACG-3′ |
| Sptk3_CPER_fragment_4_syn_F (*mcp*) | 5′-ACTTCAACCTGGTCGTGAcGC-3′ |
| Sptk3_CPER_fragment_3_syn_R (*mcp*) | 5′-GGAATGCACGGAATGCgTCACG-3′ |
| Sptk3_CPER_fragment_4_fs_F (*mcp*) | 5′-ACTTCAACCTGGTCGTGAGC-3′ |
| Sptk3_CPER_fragment_3_fs_R (*mcp*) | 5′-GGAATGCACGGAATGCTCACG-3′ |
| Sptk3_CPER_fragment_1_stop_F (*R1*) | 5′-TGGAACGCCAATGAGAtgaGG-3′ |
| Sptk3_CPER_fragment_4_stop_R (*R1*) | 5′-CACCACAACAACCtcaTCTCATTG-3′ |
| Sptk3_CPER_fragment_1_syn_F (*R1*) | 5′-TGGAACGCCAATGAGAGGcGG-3′ |
| Sptk3_CPER_fragment_4_syn_R (*R1*) | 5′-CACCACAACAACCgCCTCTCATTG-3′ |
| Sptk3_R1_del_R | 5′-tctttgtaagaaagctacccaaggCCACCTTTAGTACCAATTCCACC-3′ |
| Sptk3_R1_del_F | 5′-ggtggaattggtactaaaggtggCCTTGGGTAGCTTTCTTACAAAGA-3′ |

**Amplification of PCR fragments**

Each fragment used for CPER was amplified from Sputnik genomic DNA by PCR using 2× KOD One PCR Master Mix (TOYOBO) and 0.3 μM of each primer. PCR was performed as follows: 45 cycles of 98°C for 10 s, 55°C for 5 s, and 68°C for 32 s.

The PCR products were separated by electrophoresis on a 1% agarose gel. Bands corresponding to the expected fragment sizes were excised and purified using the Monarch Spin DNA Gel Extraction Kit (NEB, T1120S). Fragment concentrations were measured using a DeNovix DS-11 spectrophotometer (SCRUM Inc.), and molar concentrations were calculated using dsDNA (mol) = dsDNA (g) / (Length(bp) × 615.96 g/mol/bp + 36.04 g/mol).

For deletion mutants, the upstream and downstream region segments were amplified and purified as above. The two segments were subjected to an overlap extension PCR without primers to generate a fused deletion-containing template by using 2× KOD One PCR Master Mix. The reaction was performed as follows: 10 cycles of 98°C for 10 s, 60°C for 5 s, and 68°C for 35 s. The PCR product was further PCR-amplified by adding 0.3 µM of the outer forward and reverse primers. The PCR reaction was performed as follows: 35 cycles of 98°C for 10 s, 55°C for 5 s, and 68°C for 32 s.

**Assembly of Sputnik genomic DNA**

CPER was performed in a 50 µL reaction using 2.5 U of PrimeSTAR GXL DNA Polymerase with 200 µM of each dNTP and approximately 0.01 pmol of each CPER fragment. The reaction was performed as follows: initial denaturation at 98°C for 30 s, followed by 30 cycles of denaturation at 98°C for 10 s, annealing at 60°C for 20 s, and extension at 68°C for 10 min. The final extension was performed at 68°C for 10 min. The resultant CPER products were stored at 4°C until transfection.

**Genotyping Sputnik virophages.**

P1 supernatants were collected as described above and filtered through a 1.2 µm filter. One mL of the supernatant was further filtered through a 0.22-µm filter, and DNA was extracted using NaOH as described above. Regions including the mutated site were PCR-amplified using 2× KOD One PCR Master Mix and 0.3 µM of each primer as follows: 45 cycles of 98°C for 10 s, 45°C for 5 s, and 68°C for 5 s. The amplicon was purified as described above and subjected to Sanger sequencing with the PCR primers. The primers are listed below.

| | |
|---|---|
| Sptk3_mcp_Genotyping_F | 5′-TGGACAGATCAAACTGGTGG-3′ |
| Sptk3_mcp_Genotyping_R | 5′-TGCCATTTGAGTAGGATCAGC-3′ |
| Sptk3_R1_Genotyping_F | 5′-TTGTCACAAACCGTAACAGTATC-3′ |
| Sptk3_R1_Genotyping_R | 5′-CCAGGATCATCTTGTTCTTCATCTG-3′ |
